# Activated Sludge Microbial Community Assembly: The Role of Influent Microbial Community Immigration

**DOI:** 10.1101/2023.01.25.525574

**Authors:** Claire Gibson, Shameem Jauffur, Bing Guo, Dominic Frigon

**Author notes:** Address correspondence to Dominic Frigon.

## Abstract

Wastewater treatment plants (WWTPs) are host to diverse microbial communities and receive a constant influx of microbes from influent wastewater, however the impact of immigrants on the structure and activities of the activated sludge (AS) microbial community remains unclear. To gain insight on this phenomenon known as perpetual community coalescence, the current study utilised controlled manipulative experiments that decoupled the influent wastewater composition from the microbial populations to reveal the fundamental mechanisms involved in immigration between sewers and AS-WWTP. The immigration dynamics of heterotrophs were analysed by harvesting wastewater biomass solids from 3 different sewer systems and adding to synthetic wastewater. Immigrating influent populations were observed to contribute up to 25 % of the sequencing reads in the AS. By modelling the net growth rate of taxa, it was revealed that immigrants primarily exhibited low or negative net growth rates. By developing a protocol to reproducibly grow AS-WWTP communities in the lab, we have laid down the foundational principals for the testing of operational factors creating community variations with low noise and appropriate replication. Understanding the processes that drive microbial community diversity and assembly is a key question in microbial ecology. In the future, this knowledge can be used to manipulate the structure of microbial communities and improve system performance in WWTPs.

**Importance:** In biological wastewater treatment processes, the microbial community composition is essential in the performance and stability of the system. To allow future process optimisation to meet new treatment goals, we need a better understanding of factors influencing the microbial community assembly in WWTPs. This study developed a reproducible protocol to investigates the impact of influent immigration (or perpetual coalescence of the sewer and activated sludge communities) with appropriate reproducibility and controls. We demonstrate herein that influent immigration contributed up to 25 % of the sequencing reads in the activated sludge under the studied conditions, highlighting the need to consider this process in future WWTP modelling and design.

## Introduction

In activated sludge wastewater treatment plants (AS-WWTPs), the microbial community composition is intricately related to the performance and stability of the system. However, a comprehensive understanding of the structuring process of the community remains elusive. During the activated sludge (AS) treatment process, a diverse heterotrophic microbial community is grown in the plant to become highly effective at degrading the organic pollutants contained in the incoming wastewater. In recent years, efforts have been made to modify these processes to improve performance and meet new objectives for wastewater management. Modifications often require control over specific functional populations within the diverse AS microbial community. For example, enhanced phosphate removal processes require the growth of Polyphosphate Accumulating Organisms (e.g., members of the genera *Candidatus* Accumulibacter and *Tetrasphera*) (1), whilst to curtail operational problems a reduction in the abundance of detrimental bulking and foaming bacteria (e.g., genera *Gordonia* and *Thauera*) is desired (2). With a limited knowledge of community assembly mechanisms, modification and optimization of the AS process is challenging. A greater understanding of the underlying microbial mechanisms involved in the community assembly and maintenance is crucial to benefit the engineering and function of these systems.

Previous focus on microbial community assembly has been placed on the impact of temporal differences (3), wastewater characteristics (4), operational conditions (5, 6) and geographical location (7) on the AS community. However, the role of the diverse and continuously immigrating influent microbial community is poorly understood. This is a special case of community coalescence with a perpetual occurrence which likely drive in part the community dynamics (8).

The community in municipal wastewater originates from numerous environments including human faeces, sewer biofilm and sediments, and soil runoffs (3). Studies have produced conflicting results on the relative importance of influent immigration on the assembly of the AS community. Some suggested that the overall effect of the influent community is negligible, as they observed that the AS microbial community remains stable over time despite changes in the composition of the influent wastewater community (4). The activity of immigrating taxa has also been questioned, with some studies showing these populations often have low or negative net growth rates (9, 10).

Other literature reports argued influent immigration to be an important process in microbial community assembly. In lab-scale AS reactors operated at extremely low temperatures and solids retention times, immigration from sewers was found to be essential to maintain complete nitrification (11). The occurrence of shared operational taxonomic units (OTUs) between influent and AS communities also infers high immigration rates in full-scale AS-WWTPs (9, 12). Furthermore, a recent study of 11 AS treatment plants in Denmark, selected based on their highly similar process design, identified the AS community composition to be strongly influenced by the influent wastewater community composition (13). However, it remains difficult to ascertain if the influent community composition is a factor independent from the substrate composition of the wastewater, which precludes strong conclusions.

With contrasting reports on the impact of immigration, it is challenging to understand the relative importance of this process. This variability can be somewhat explained by the limited methods used to study the immigration process, which lack reproducibility and assessment of the activity of immigrants. As reviewed by Mei & Lie (2019), commonly used approaches include counting shared microbial species between ecosystems (i.e., the influent and AS-WWTP), microbial source tracking and neutral community modelling (14). However, these approaches only imply a contribution from the immigrating source and provide little information about the fate and activity of these organisms. Furthermore, neutral modelling approaches to investigate immigration (such as the neutral theory of biodiversity) assume that organisms are functionally equivalent (15), this assumption is likely inaccurate considering the variety of substrates available to the heterotrophic populations (16) and the niche differences between the influent and AS. It has been proposed that immigrants should be classified as either rare diffusive immigrants, or time continuous high-rate mass flow immigrants (17). Heterotrophic mass flow immigrants appear to be heavily influenced by deterministic selection, suggesting that they should be divided into relevant functional guilds when assessing their assembly mechanisms. Consequently, we cannot rely on mathematical modelling approaches alone to resolve the relative importance of selection, immigration, and drifts in the assembly of the community. Furthermore, full-scale systems display too much variability in terms of wastewater composition, operation and community to allow appropriate reproducibility and provide an accurate assessment of the mechanisms at play during immigration.

Given the complex behaviour of microbial communities within wastewater, reproducible and highly controlled systems are required to accurately investigate and quantify the impact of immigration into AS-WWTPs and determine the fate of immigrating bacteria. Because the presence of specialized populations in the influent wastewater is often correlated with the presence of specific substrates (18; Chapter 7), it is impossible to understand immigration independently from the wastewater substrate compositions in full-scale systems. Therefore, here we propose the development of a system where the substrate landscape and the immigrating community can be manipulated independently using highly controlled laboratory-scale reactors, to investigate fundamental questions on the quantitative impact of immigration and activity. To address the limitations observed in past immigration studies, this work aimed to decouple the influent substrates and microbial community, and specifically investigate the impact of immigration with appropriate reproducibility and controls. Using a series of highly controlled reactors (a total of 72 reactors were operated for 17-25 weeks), we manipulated the influent microbial community independently to identify the effect of constant immigration on the AS microbial community. The laboratory-scale AS reactors were inoculated with mixed liquors from three different AS-WWTPs, representing three of the most different municipal AS microbial communities among the AS-WWTPs located within 100 km of Montréal (Québec, Canada). The purpose of this was to determine whether the impact of immigration was dependent on the established community within the reactors. This also evaluated the reactor protocol for future use with different but similar starting communities. Finally, dynamics in the AS microbial community compositions were analysed by 16S rRNA gene amplicon sequencing. The long-term goal of the current work is to provide a reproducible protocol to conduct studies on the fate of immigrants and their activities in the AS mixed liquor microbial communities, and to develop an appropriate method to study this aspect of biological wastewater treatment systems.

## Material and Methods

### Experimental Design and Reactor Setup

In total, 3 ***sets*** of reactors were operated (Set A, B and C), which differed based upon the source of influent solids used. Figure 1b shows a basic schematic of *one* set of reactors operated. For the first set of reactors (Set A), the reactors received La Prairie (Québec, Canada) influent solids and were operated from May until September 2016. For the second and third sets (Set B and Set C), the test reactors received Cowansville and Pincourt (Québec, Canada) influent solids, respectively, and were operated from May until November 2017. To investigate the impact of the starting community, each ***set*** of reactors were divided into 3 ***blocks*** (Block a, b and c), the reactors in each block were inoculated with mixed liquor obtained from either La Prairie, Cowansville or Pincourt AS-WWTPs, respectively (Figure 1b).

**Figure 1.**
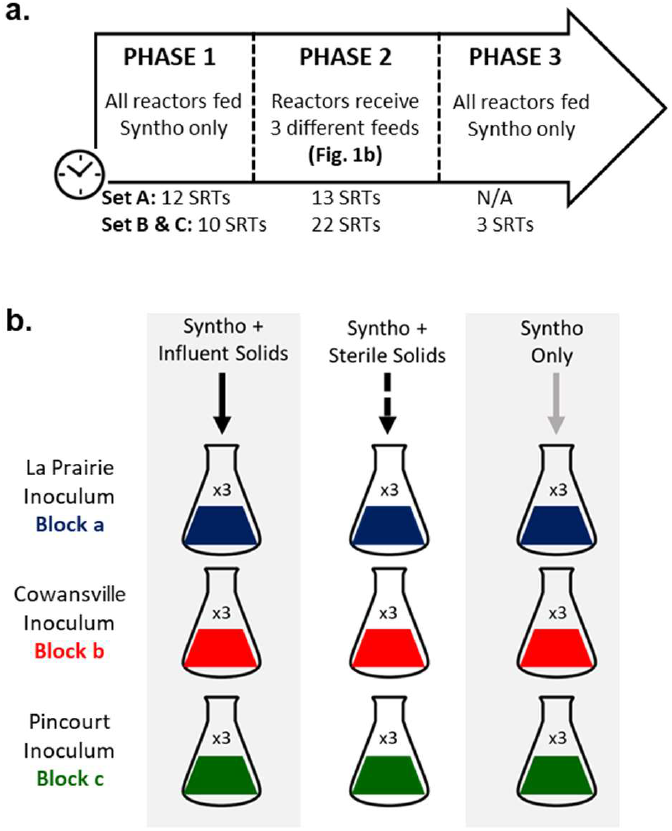
Experimental set up. (a) Schematic of one set of reactors (i.e., receiving one source of influent solids) in Phase 2. Three sets of reactors were operated in total. Set A received La Prairie influent solids; Set B, Cowansville influent solids; Set C, Pincourt influent solids. The block describes the inoculum mixed liquor source. (b) Timeline of reactor operation. Reactors were operated in three phases with feed altered.

The three wastewater treatment plants were selected as a source of influent solids and mixed liquor inocula from among 8 AS-WWTPs within a diameter of approximately 100 km from Montréal (Québec, Canada) based on the results of a previous study. This study revealed that the compositions of the activated sludge mixed liquor microbial communities at the three selected plants were at the extremes of the composition distribution when visualized using a principal coordinate analysis plot in a previous study (19). In other words, the mixed liquor communities showed the greatest differences among the sites tested.

The reactors were operated in three phases (Figure 1a), with the feeds altered to investigate the effect of immigration. During Phase 1, all reactors received Syntho (see section *Feed Preparations*) only to develop a steady-state core microbial community adapted to the specific substrate composition. At the end of Phase 1, the mixed liquors of all 9 reactors that had received the same inoculum (i.e., 1 block of reactors; Figure 1b) were combined and split again to ensure a homogeneous stable community in each reactor as described by Kaewpipat and Grady, 2002 (20). During Phase 2, the 3 test reactors in each inoculum block received Syntho supplemented with influent solids (resulting in 9 test reactors per set) to simulate immigration into the AS. In addition, nine reactors (3 per block) were defined as substrate controls and received Syntho supplemented with sterile influent solids (autoclaved at 121°C, 15 psi for 30 minutes). The final 9 reactors (3 per block) received Syntho only to act as a continuity control (9 per reactor set). In total, 72 reactors were operated comprising of 3 blocks of inoculum and 3 sources of influent solids, operated in triplicate resulting in 27 reactors per influent solids used. In 2017, there was only one set of Syntho only controls used because Sets B (Cowansville influent solids) and Set C (Pincourt influent solids) were operated simultaneously and their Syntho only control reactors were equivalent (which allowed the economy of 9 reactors). A detailed overview of the reactor experimental design is outlined in Table S2.

### Reactor Operation

The reactors were inoculated with 90 mL of mixed liquor (2.7 g-VSS/L) obtained from La Prairie (block a), Cowansville (block b) and Pincourt (block c) AS-WWTPs. The AS mixed liquor (ML) samples from the 3 AS-WWTPs were collected within a period of 4 hours (block a; May 2016, block b and c; May 2017). Samples were transported on ice, stored at 4 *°C* and processed within 24 hours of collection. All reactors were incubated at 21 °C with shaking at 180 rpm to allow aeration and gentle mixing. The average reactor hydraulic retention time (HRT) was 1.8 days, and the solid retention time (SRT; the average time solids were maintained within the reactors) was 5 days. The HRT and SRT were controlled through daily feeding and wasting. Three times a week, the total reactor volume was settled in a 100-mL graduated cylinder for 45 min to allow solid-liquid separation to mimic the clarifier of an activated sludge wastewater treatment process, and the appropriate volume of the supernatant was removed.

### Influent Solid Processing for Phase 2 Feed

Influent wastewater solids were concentrated at the AS-WWTP by settling influent wastewater for 15 min in 20-L buckets. Approximately 10-L of concentrated influent wastewater was transported to the lab on ice and processed immediately. To separate the influent wastewater and solid fractions, the wastewater was centrifuged in 50 mL aliquots at 21,100 × g for 10 min (Thermofisher Scientific model ST16R Centrifuge). The supernatant was removed, and the solids were re-suspended in Synthetic wastewater (Syntho; defined in section Feed Preparations) to wash. The centrifugation and wash step were repeated three times, and the influent solids were resuspended in a final ~800 mL of synthetic wastewater. The volatile suspended solids (VSS) concentration of the stock influent solid solution was measured using the Standard Method 2540E (21) before dilution. The concentrated influent solids stock was stored at 4 *°C* for no more than two weeks before use.

### Feed Preparation

A stock solution of influent solids was prepared as described in section *Influent Solid Processing for Phase 2 Feed*. Synthetic wastewater ‘Syntho’ was produced using a modified recipe as described by Boeije et al., 1999 (22), and autoclaved in 500 mL containers at 121 *°C* and 15 psi for 30 minutes. The ‘Syntho + Influent Solids’ feed was prepared by diluting the stock influent solid solution in Syntho to a concentration of 120 mg-VSS/L, which represents the average VSS concentration of influent wastewater reported in the three wastewater treatment plants studied (23). The ‘Syntho + autoclaved solids’ feed was prepared in the same way with the addition of autoclaving at 121 °C/15 psi for 1 hour to sterilise.

### Sample Analysis

Mixed liquor reactor samples were collected weekly for analysis of total suspended solids (TSS) and volatile suspended solids (VSS) using Standard Methods 2540B and 2540E respectively (21). Effluent wastewater (the supernatant after settling) samples were collected for the analysis of chemical oxygen demand (COD) using Standard Methods 5220D (21). Biomass samples for DNA extraction were centrifuged and stored ready for nucleic acid extraction at −80 *°C* until further use.

### 16S rRNA Gene Amplicon Sequencing

DNA was extracted from stored biomass samples using DNeasy PowerSoil Kit (Qiagen, Germantown, MD, USA). PCR of the 16S rRNA gene V4 region was conducted using 515F (5’-GTGYCAGCMGCCGCGGTAA-3’) and 806R (5’-GGACTACNVGGGTWTCTAAT-3’) primers (24, 25). PCR conditions were as follows: 94 °*C* for 3 min followed by 35 cycles of 94 °*C* for 45 sec, 50 °*C* for 60 sec, 72 °*C* for 90 sec. 72 °*C* for 10 min and a final hold at 4 °*C*. Unique Uniprimer barcodes were added to the amplicons from each sample in a second PCR reaction to allow sample pooling. Reaction conditions were as follows; 94 °*C* for 3 min followed by 15 cycles of 94 °*C* for 30 sec, 59 °*C* for 20 sec, 68 °*C* for 45 sec. 68 °*C* for 5 min and a final hold at 4 °*C*. Barcoded amplicons were purified using QIAquick PCR Purification Kit (Qiagen, Germantown, MD, USA) and pooled at equimolar concentrations. Amplicons were sequenced on the Illumina MiSeq PE250 platform at McGill University and Génome Québec Innovation Centre (Montréal, QC, Canada).

### Bioinformatic and Statistical Analyses

Sequencing data was analysed using QIIME 2 (26) and R Software (packages “vegan” and “ape”). Specifically, the raw sequences were denoised and errors corrected using DADA2 (27) in Qiime2 pipelines. Sequences were rarefied to 40,000 reads in R. ASV tables were exported directly from Qiime2 after quality filtering. Taxonomy was assigned using MiDAS 2.0 reference database (28) and tabulated. Microbial community diversity was analysed using the “vegan” package of R (29) and Jaccard dissimilarity.

Analysis of Similarities (ANOSIM) was conducted using the using the “vegan” package of R (30) to statistically test the significance of differences between groups. The ANOSIM test compared the mean of ranked dissimilarities between groups with the mean of ranked dissimilarity within groups. An ANOSIM R value close to ‘ 1’ indicates dissimilarity between groups, whilst a value close to ‘0’ indicated that the similarities between groups and within groups were on average the same (30).

An odds ratio test was used to assess whether the growth rate of immigrants impacted their fate during Phase 3 of reactor operation. Values were calculated by dividing the odds of the first group (genera with a positive net growth rate persisting at the end of Phase 3) by the odds of the second group (genera with a negative net growth rate persisting at the end of Phase 3).

### Definitions of Population Categories

The **core *resident populations*** were present in the majority of reactors and followed the criteria outlined by Wu et. al 2019 for core communities (7). Taxa were filtered based on the overall abundant genera, by calculating the mean relative abundance of a given genera. Genera with a mean relative abundance of over 0.1% were selected as overall abundant taxa. Ubiquitous genera between reactors were identified based on their occurrence in over 80% of the reactor set (where n=27; genera must have occurred in at least 22 reactors). Taxa that were present in reactors receiving either Syntho only or Syntho + influent solids but that appeared more sporadically (i.e., in fewer than 80% of reactors) or in lower abundance (below 0.1 %) were classified as the ***non-core resident populations***.

The populations of immigrating bacteria were determined and classified by comparing the reactors receiving influent solids with those receiving Syntho only (i.e., without active immigration). Bacteria present only in reactors receiving Syntho + influent solids (or sterile influent solids) were classified as influent immigrants, the presence of which was dependent upon immigration. Taxa occurring in both the reactors receiving influent solids and sterile influent solids were filtered based upon abundance; those occurring in equal abundance under both conditions were considered to be ***residual immigrants*** or residual DNA, whilst those occurring in higher abundance in the reactor receiving active solids were classified as ***growing immigrants*** (or ***actively growing immigration-dependent populations***).

### Quantifying Net Growth Rate with Steady-State Modelling

The impact of immigration on the average growth of taxa in activated sludge systems can be visualized using a scatter plot of their relative abundances in the mixed liquor vs. the influent wastewater. A steady-state mass balance on biomass and substrates can be used to develop a model quantifying the levels of immigration and the growth rate and mapping them on the relative abundance scatter plot. The model used here was partially developed by Grady et al., 2011 (31) and it is an extension of the model presented by Mei et al., 2019 (14). A detailed development is available in Guo, 2022 and its key elements are presented here for convenience (32).

With a mass balance on the biomass of the i-th taxon for an activated sludge system assuming completely mixed biomass, it is possible to show that the i-th taxon specific growth rate (*μ_i_*) is given by eq 1.

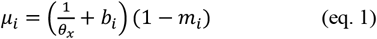

Where *θ_x_* is the solid retention time, *b_i_* is the decay rate of the i-th taxon, and *m_i_* is defined as the immigration level of the i-th taxon defined by eq. 2.

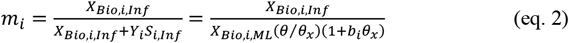

Where *X_Bio,i,inf_* and *X_Bio,i,ML_* are the biomass concentrations of the i-th taxa in the influent and the mixed liquor, respectively, *S_i,Inf_* is the concentration of substrates consumed by the i-th taxon, *y_i_* is the biomass yield of the i-th taxon on the substrate consumed, and *θ* is the hydraulic retention time.

To map the growth rates and immigration levels of the relative abundance scatter plots, eq. 1 and eq. 2 can be developed considering the capture of the influent biomass of the i-th taxon by the mixed liquor solids (*f_OHO,Capt_*), the DNA extraction yields for the influent and mixed liquor (*γ_DNA,Inf_*, *γ_DNA,ML_*) and the total solids of the influent and the mixed liquor (*X_Tot,Inf_*,*X_Tot,ML_*). With these considerations, the relationship between the proportions of the i-th taxon in the influent and mixed liquor as determined by 16S rRNA gene amplicon sequencing (*f_16S,Inf,i_*, *f_16S,ML,i_*) is given by eq. 3.

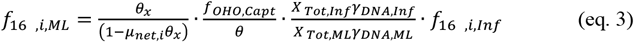

Where *μ_net,i_* is the net specific growth rate of the i-th taxon defined as specific growth rate minus specific decay rate (*μ_net,i_* = *μ_i_ - b_i_*)

The log-log version of eq. 3 (eq. 4) shows that taxa appearing on the same 45°-line (i.e., 1: 1 line) in the log-log scatter plot of relative abundances have the same net growth rate and the same immigration level, and that the y-intercept (bold term between square brackets) of this 45°-line is a function of *μ_net,i_*.

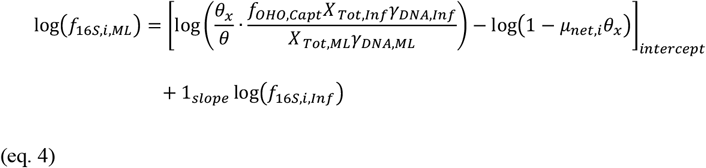

## Results

### Variations in Overall Community Compositions During Experiment

Determining the specific impact of influent immigration on the structure of the AS community in full-scale systems presents a significant challenge because of the numerous spatial and temporal variations between locations. The current study utilised controlled reactor experiments to investigate the impact of immigration on a naturally assembled, diverse microbial community. During Phase 1, all reactors received Syntho (a synthetic wastewater; 22) to develop a steady-state core microbial community adapted to a specific substrate composition. This procedure also minimized the biomass of *residual immigrants* (immigrant populations not consuming resources as defined in section Definitions of Population Categories) from the mixed liquor inocula remaining in the system prior to the test period (i.e., Phase 2).

During Phase 1 when fed with Syntho only, the microbial communities of the reactors became more similar to one another when compared to the differences among starting inocula (Figure 2), and the amplicon sequencing variant (ASV) richness decreased (Figure S1). Irrespective of the inoculum, 46.2 + 4.7 % (+ indicates standard deviation) of the observed genera at the end of Phase 1 were shared between the 27 resulting AS reactor communities forming the *core resident populations* of Syntho (as defined in section Definitions of Population Categories), whilst at the beginning of Phase 1, only 27.1 + 4.7 % of the observed genera were shared among the 3 inocula. Nonetheless, the majority of AS reactor communities at the end of Phase 1 remained significantly clustered based on the inoculum received according to both Jaccard and Bray-Curtis dissimilarities. The ANalysis Of SIMilarity (ANOSIM) can be used to express in a common quantity the differences between clusters, with ANOSIM R values closer to 1.0 indicative of higher dissimilarity between groups, whilst those closer to 0.0 indicating higher similarity. Comparing Jaccard and Bray-Curtis dissimilarities among communities at the end of Phase 1 revealed a higher difference measured by Jaccard dissimilarities (ANOSIM R = 0.88 + 0.05) than Bray-Curtis dissimilarity (ANOSIM R = 0.61 + 0.04; Figure 2). The Jaccard dissimilarity is more sensitive to the least abundant populations than the Bray-Curtis dissimilarity, this suggests that the main difference between the communities assembled on Syntho during Phase 1 was in low abundance populations. These observations show that Syntho selected for a *core resident populations* comprising of the most abundant populations. Whilst populations specific to the inoculum community were in lower abundance. As the experiment progressed, the microbial community of most reactors receiving the same feed remained clustered based on the inoculum (Table S3) with the exception of a few outliers as visualised in Figure S1.

**Figure 2.**
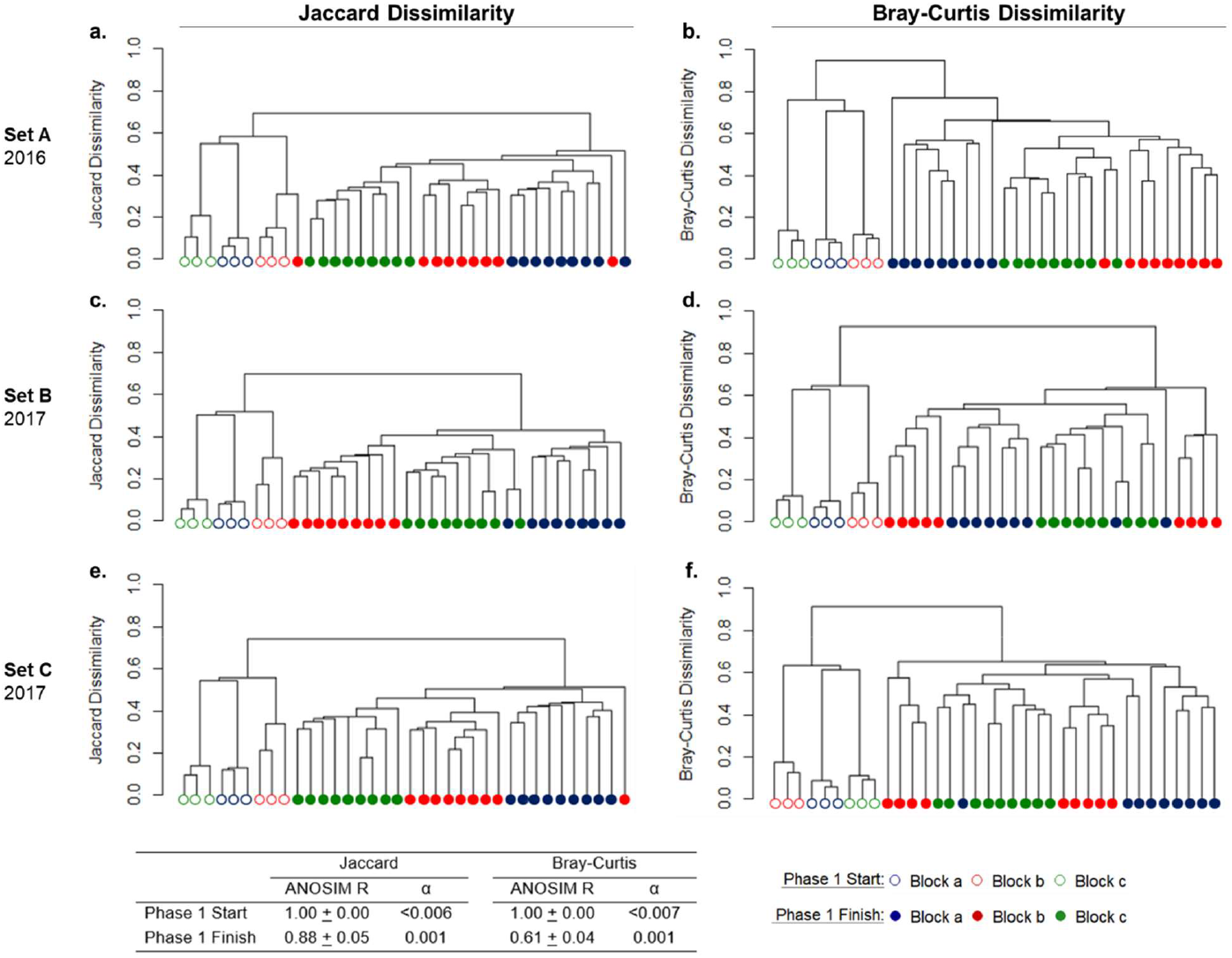
Tree dendrogram of microbial community at the start (hollow symbols) and end (filled symbols) of Phase 1 using UPGMA clustering method and the Jaccard dissimilarity (a, c, e) or the Bray-Curtis dissimilarity (b, d, f). Set A (a, b) were operated in 2016, whilst Sets B (c, d) and Set C (e, f) were operated in 2017. The Block indicates the source of inoculum: Block a-La Prairie mixed liquor, Block b-Cowansville mixed liquor, Block c-Pincourt mixed liquor. Inoculum communities were the same for Set B and C as they were operated in tandem. Each inoculum (hollow symbols) represents a single biological sample that was extracted and sequenced 3 times, whilst the end of Phase 1 (filled symbols) are communities raised in independent reactors.

During Phase 2, reactors received three different feeds. The test reactors received a feed comprising of Syntho supplemented with influent solids to determine the impact of immigration. The remaining reactors received Syntho and sterile solids or Syntho only, which acted as substrate and continuity controls, respectively. At the end of Phase 2, principal coordinate analysis using Jaccard dissimilarity (Figure 3) revealed that immigration caused the resulting communities to become more similar to the influent community, as shown by the points representing these communities being located closer to the influent community along the PCoA1 axis. The microbial community of the reactors receiving sterile solids also moved away from the starting position, likely due to the additional substrate and residual DNA in the autoclaved influent biomass (Figure 3).

**Figure 3.**
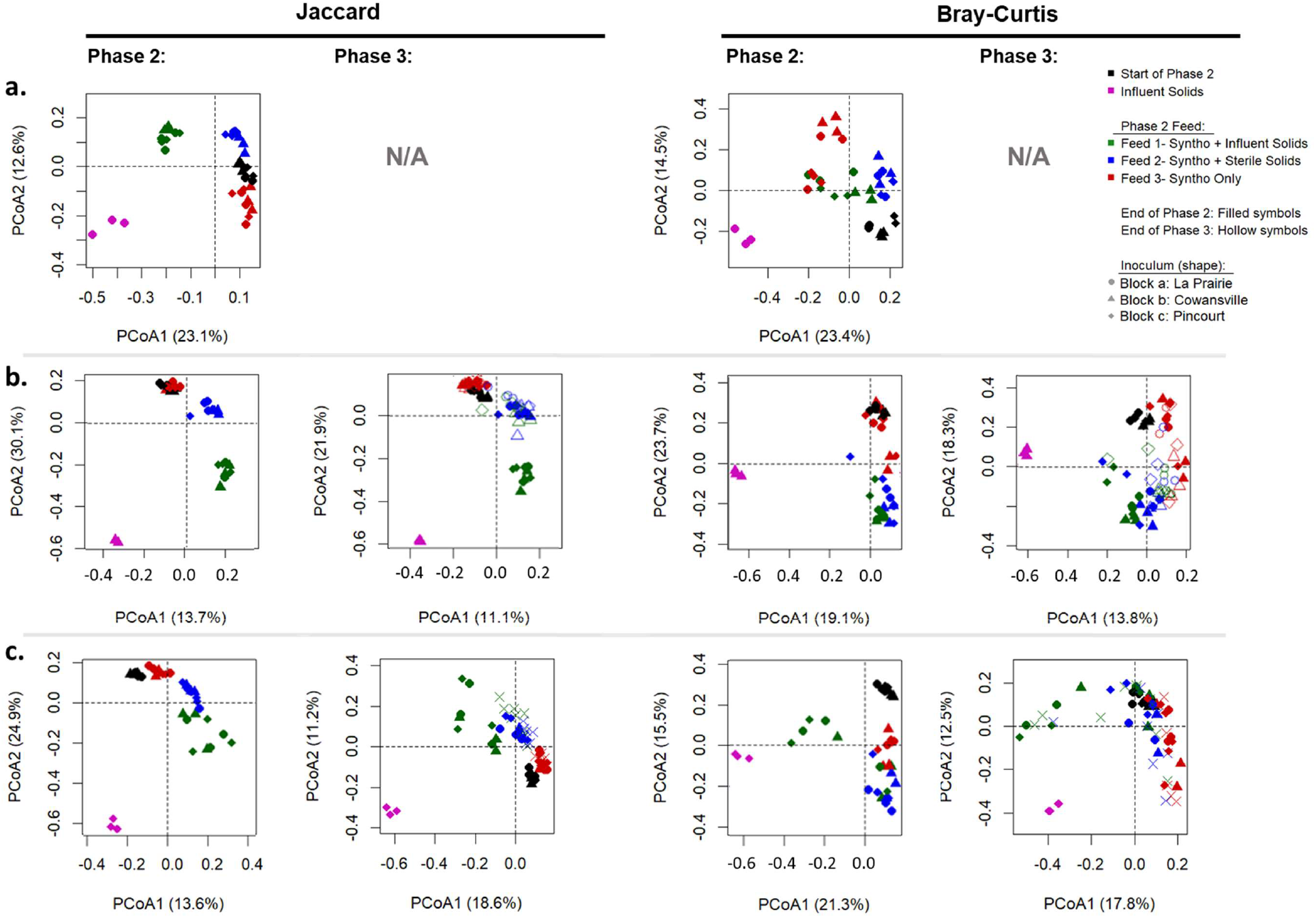
Principal coordinate analysis of reactors communities at the end of Phase 2 and 3, visualised at ASV level using Jaccard and Bray-Curtis Dissimilarity. a) Set A-La Prairie Influent Solids b) Set B-Cowansville influent solids c) Set C-Pincourt influent solids. Filled symbols represent samples taken at the end of Phase 2 and hollow symbols were samples collected at the end of Phase 3.

Unlike with Jaccard dissimilarity, when visualising the community composition data using Bray-Curtis dissimilarity (Figure 3), the impact of immigration on the overall community was less apparent, particularly in reactors of Set A and B. In these reactors, the change in the microbial community during Phase 2 was similar in the reactors with immigration and the sterile controls which received sterilised influent solids. Given the higher sensitivity of the Jaccard dissimilarity than the Bray-Curtis dissimilarity for the presence of low abundance population, it appears that immigration had more of an impact on these populations that are likely maintained by their continuous influx in the mixed liquor from the influent.

During Phase 2, the ASV richness of reactor Set A (test reactors receiving La Prairie influent solids) and B (test reactors receiving Cowansville solids) increased (Figure S2). This increase was not observed among the flasks receiving sterile influent solids nor Syntho only throughout, thus indicating that the immigrant populations are associated with the changes. The impact of immigration on the richness of reactor Set C was more variable. The original richness of the reactors was not restored to that of the starting community (Figure S2), likely due to the relative substrate-composition simplicity of Syntho compared to actual wastewater and the greater homogeneity of laboratory-scale flask reactors than of full-scale AS-WWTPs.

Phase 3 of reactor operation was introduced to determine whether the impact observed with immigration could be sustained over time without continuous seeding of influent solids. During this phase, influent solids were removed from feeding, and all reactors received Syntho only as in Phase 1. Within 3 SRTs, the communities in reactors which previously received active influent solids during Phase 2, moved back towards the starting position (start Phase 2) and became more similar to the communities in reactors which received sterile inactive influent solids throughout (average Jaccard distance between reactors with immigration and sterile controls reduced from 0.56 to 0.47 in Set B and 0.59 to 0.49 in Set C indicating increased similarity; Figure 3) demonstrating that the full impact of immigration could not be maintained over time without continuous seeding.

### Classification of Genera

To provide a more detailed analysis of the impact of immigrating taxa, the genera forming the microbial community at the end of Phase 2 were classified in four categories according to their presence in reactors receiving influent solids. This analysis was conducted at genus level as ASVs within the same genus are likely to have similar ecological functions, and because direct comparisons of ASVs between the influents and the mixed liquors were often unreliable due to their low abundances in one of the two compartments. The four categories assigned are outlined in the Definitions of Population Categories section and are briefly recalled here. First, a genus was defined as a ***core resident population*** if it occurred in at least 80 % of all the reactors within a set with a relative abundance of at least 0.1 %. Second, ***non-core resident genera*** appeared in reactor without immigration, but in fewer than 80 % of these reactors or at an average abundance below 0.1 %. Third, ***growing immigrant genera*** were only present in the reactors receiving influent solids and were at a higher abundance than in the reactors receiving autoclaved solids in the same set (i.e., same influent solids source). These genera were presumed to be actively growing. Fourth, ***residual immigrant genera*** were those present only in the reactors receiving influent solids and occurring in the reactors with immigration in equal or lower abundance to the sterile autoclaved control group within the same set.

The resident community in all reactors consisted of between 50 and 73 genera depending on the set of reactors (i.e., influent community), and these genera accounted for between 69 and 76 % of the total reads observed in the reactor set (Table 1). Conversely, the immigrant populations showed a higher diversity (118 to 206 genera) but accounted for much fewer reads from the reactor set (4 % to 14 %). This high richness and low abundance of the immigrant populations is in line with the impact of immigration being more visible when using the Jaccard dissimilarity, but not when using the Bray-Curtis dissimilarity (Figure 3). Therefore, operation and the Syntho substrate composition appeared to have determined the resident abundant members of the communities, whilst immigration contributed towards low abundance members of the communities.

**Table 1:** Percentage of overall reads which form the resident and immigrant communities at the end of Phase 2 for reactor sets receiving different influent solids

|  | Set A:<br>La Prairie |  | Set B:<br>Cowansville |  | Set C:<br>Pincourt |  |
| --- | --- | --- | --- | --- | --- | --- |
|  | Genera<sup>1</sup> | Reads (%)<sup>2</sup> | Genera | Reads (%) | Genera | Reads (%) |
| Core Residents | 73 | 75.3 ± 5.4 | 54 | 76.0 ± 4.9 | 50 | 69.3 ± 11.5 |
| Non-Core Residents | 133 | 15.7 ± 4.3 | 121 | 7.9 ± 1.7 | 122 | 22.4 ± 12.8 |
| Growing Immigrants | 127 | 8.2 ± 2.0 | 206 | 14.3 ± 4.9 | 118 | 4.2 ± 1.4 |
| Residual Immigrants | 7 | 0.3 ± 0.2 | 5 | 2.1 ± 1.6 | 12 | 2.9 ± 3.2 |

<sup>1</sup>Number of genera within the category
<sup>2</sup>Percentage of overall reads ± standard deviation

The immigrant population varied based upon the source of influent solids (Figure 4). In reactor Set A, which received influent solids from La Prairie wastewater treatment plant, the genus Spb280 (genus midas_g_81, family *Comamonadaceae;* 33), and an uncultured genus of the family *Synergistaceae* were the dominant immigrants. Whilst in Set B, members of the genus *Aquabacterium* (family *Comamonadaceae*) were dominant, and in Set C, the most abundant immigrant genus was *Dechloromonas* (family *Rhodocyclaceae)*. In addition to unique immigrant communities based on the source of influent solids, there also appeared to be some inoculum effect (between Blocks). Within the immigrant population of Set B reactors, those which were inoculated with La Prairie mixed liquor (Block A) were abundant in the genus SipK9 (genera *midas_g_1719 and midas_g_2835*, family *Rhodobacteraceae*; 33) which accounted for up to 11.7 % of the reads and explained some of the variability in percentage reads contributed though immigration reported in Table 2. Whilst in the other Set B reactors inoculated with Cowansville and Pincourt (Block b and c, respectively), SipK9 was not observed with the same high abundance. The function of SipK9 is currently unknown but given the abundance in the reactors may warrant further investigation.

**Figure 4.**
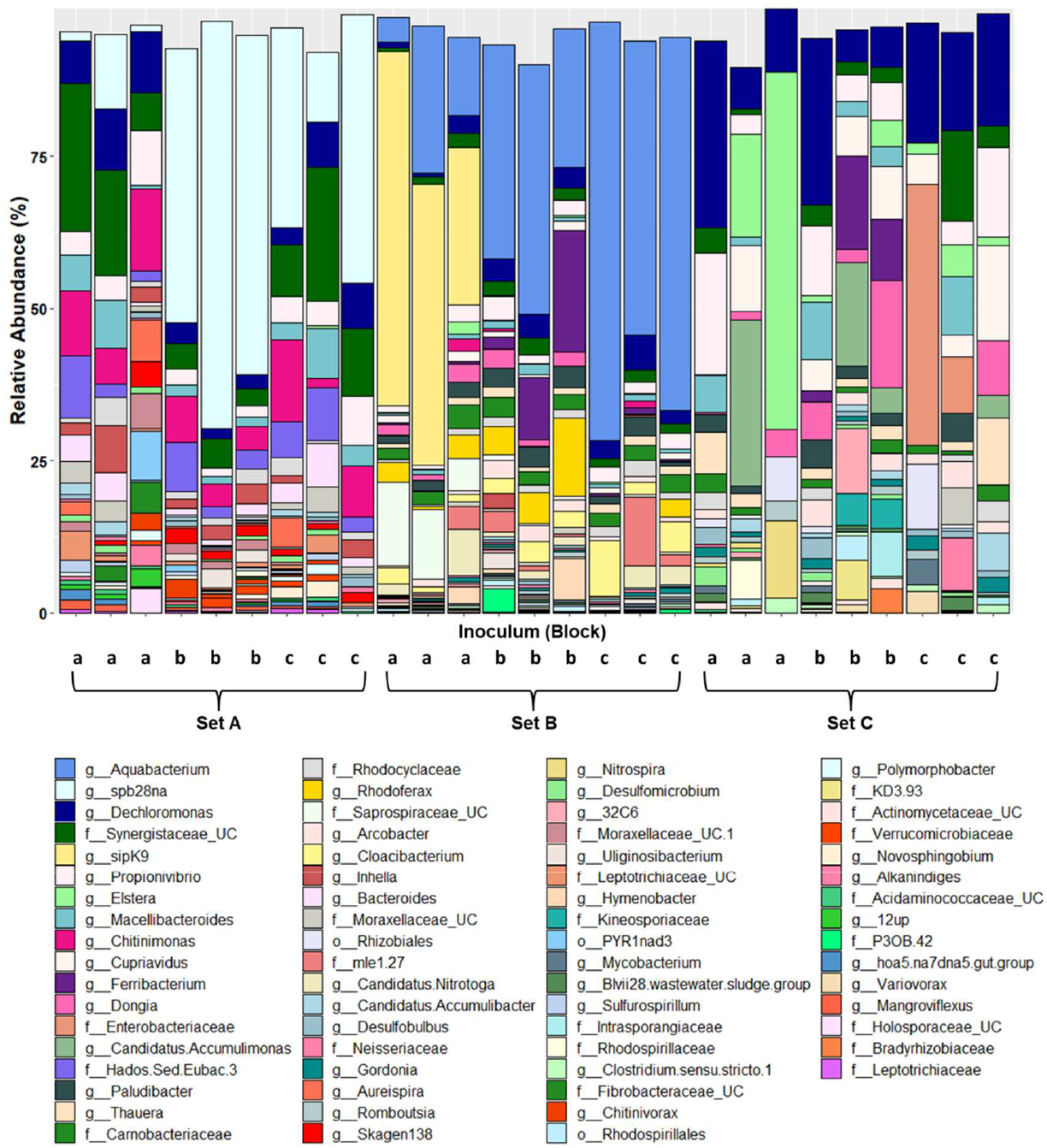
Top 70 immigrating genera, normalised to total immigrants per reactor. Inoculum A-La Prairie ML, B-Cowansville ML, C-Pincourt ML. Set A-La Prairie influent solids, B-Cowansville influent solids, C-Pincourt influent solids. UC in the legend indicates uncultured strains of a given taxa.

**Table 2:** Percentage of overall reads which form the resident and immigrant communities at the end of Phase 3

|  | Set B |  | Set C |  |
| --- | --- | --- | --- | --- |
|  | Genera | Reads (%) | Genera | Reads (%) |
| Core Residents | 54 | 85.7 ± 5.9 | 50 | 78.3 ± 15.2 |
| Non-Core Residents | 124 | 10.4 ± 5.0 | 121 | 19.4 ± 13.4 |
| Growing Immigrants | 65 | 3.5 ± 2.1 | 29 | 1.8 ± 2.7 |
| Residual Immigrants | 4 | 0.1 ± 0.1 | 9 | 0.2 ± 0.3 |

<sup>1</sup>Number of Genera within the category
<sup>2</sup>Percentage of overall reads ± Standard Deviation

During Phase 3, influent solids were removed, and all reactors received Syntho Only as in Phase 1 to determine if immigration had a lasting effect on the microbial community. Phase 3 was only conducted for reactor Set B and C in 2017. At the end of Phase 3, the number of immigrants remaining in the reactors had reduced in both reactors Set B and C (Table 2). In reactor Set C, immigrants remaining included taxa from the families *Rhodospirillaceae, Rhodocyclaceae, Nitrospiraceae* and *Flavobacteriaceae*. In reactor Set B, the families *Rhodocyclaceae, Gallionellaceae* (genus *Candidatus* Nitrotoga), *Comamonadaceae* (genus *Rhodoferax)* and *Xanthomonadaceae* (genus sipK9) were among the most abundant growing immigrants remaining. To further investigate factors influencing the persistence of immigrants in the reactors at the end of Phase 3, further analysis of the net growth rate was conducted.

### Assessing the Growth Rate of Genera

The impact of immigration on the various genera can be visualized with a log-log scatter plot of their relative abundances in the mixed liquor and in the influent microbial communities (Figure 5). Assuming that the ASV composition of each genera is the same in the influent and in the mixed liquor, the log-log scatter plot of the relative abundances of the genera can be understood as representing their net growth rate by using a mass balance model (eq. 4). Based on the model in eq. 4, genera with different relative abundances but falling on the same 45°-lines on Figure 5 have the same net growth rate. As a reference, a 45°-line was drawn on the scatter plot of Figure 5 for net growth rates (*μ_net_*) equal to 0 (i.e., the point where the growth rate of the organisms is equal to its decay rate). Any points falling above the zero net growth rate line (*μ_net_* = 0) display a positive net growth rate, whilst those below this line have an overall negative net growth rate. According to reactor theory, organisms displaying positive net growth rates should be maintained in the reactor without immigration assuming no other factors such as competition occur, whilst organisms displaying negative net growth rates would be washed out if immigration were stopped.

**Figure 5.**
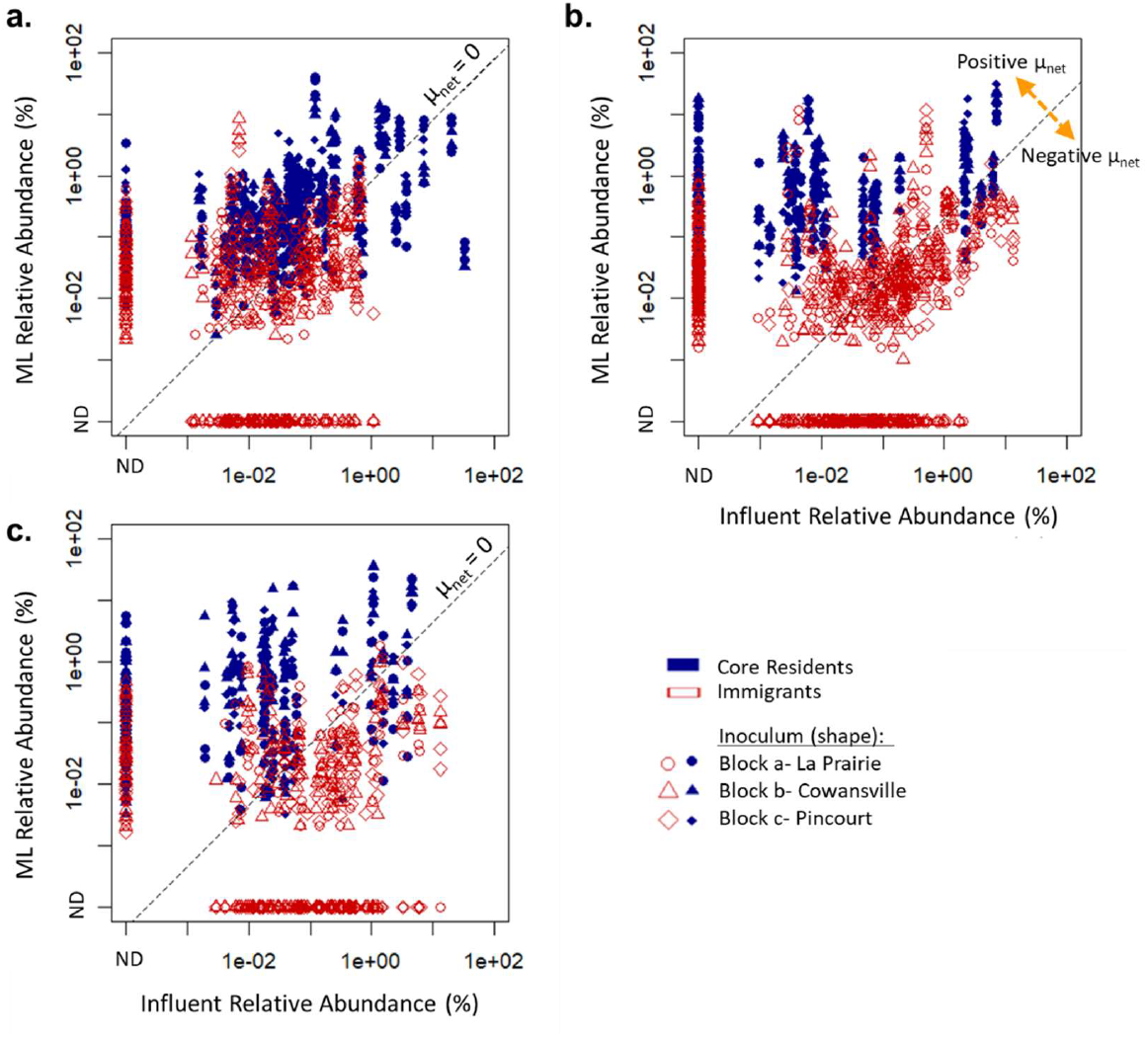
Comparison of influent and mixed liquor (ML) relative abundance of immigrant and resident genera. ‘ND’: Not Detected indicated genera that were below the detection limit. Reactors received influent solids of different sources: a) Set A-La Prairie influent solids b) Set B-Cowansville influent solids c) Set C-Pincourt. Dashed line represents a net growth rate of 0. Genera on the same 45°-lines have the same net growth rates (u_net_)

Genera classified in the core resident population by our criteria typically exhibited a positive net growth rate (Figure 5), whilst genera classified in the growing immigrant population typically had a negative net growth rate, particularly those from Cowansville and Pincourt influents (Set B and C; Figure 5b and 5c). Selection against certain genera present in the influent at high relative abundant (up to 1 % of the reads) was also observed as these genera remained undetected in the mixed liquor (Figure 5; shown as ‘ND’ in ML). Interestingly, some genera classified as immigrants were not detected (ND) in the influent, suggesting that they occurred below the detection limit in the influent solids, and increased to detectable levels within the reactor communities.

In Figure 5, it was observed that a selection of immigrants displayed a positive net growth rate. Based on reactor theory, it would be expected that these genera could be maintained within the reactors without continuous immigration. At the end of Phase 3, although reduced, a number of immigrants remained in the reactors (Table 2). Analysis of the net growth rate of these taxa (Figure 6), showed that of those remaining at the end of Phase 3 between 75 and 77 % had displayed an overall positive net growth rate during Phase 2. However, many other immigrants that had a positive net growth rate during Phase 2, were not detected at the end of Phase 3. An odds ratio test of the taxa remaining at the end of Phase 3, and their associated net growth rate evaluated in Phase 2 supported that taxa with a positive net growth rate were between 5.6 and 7.4 times more likely to persist at the end of Phase 3 than those with a negative net growth rate.

**Figure 6.**
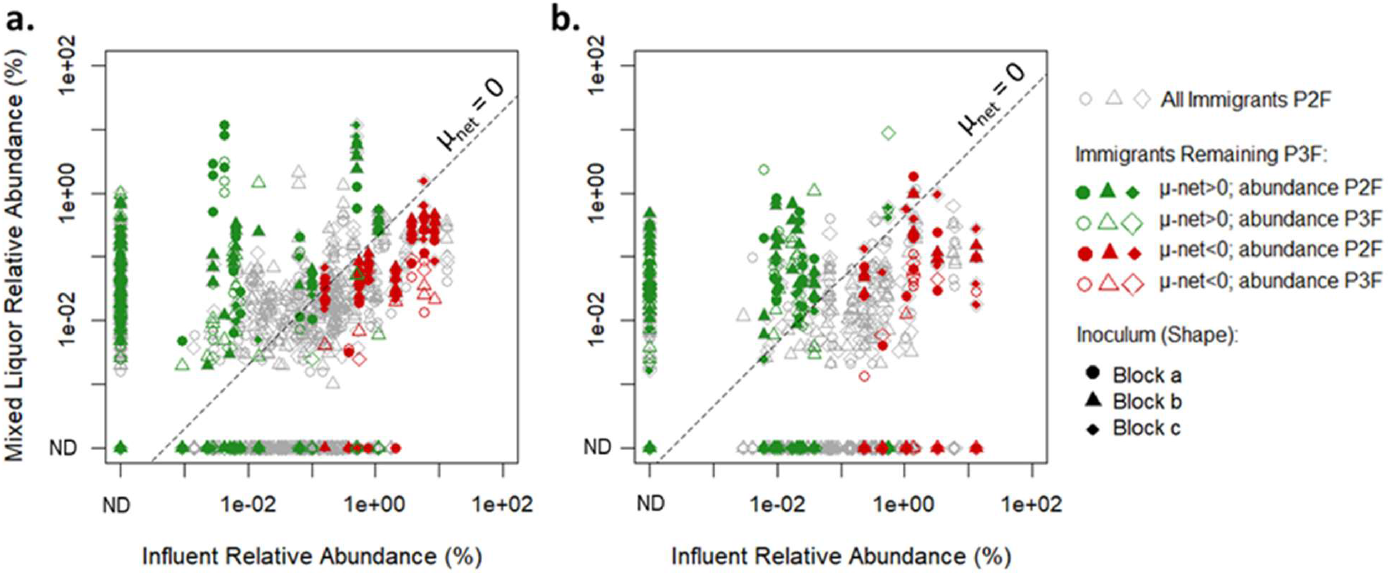
Comparison of growing immigrant genera at the end of phase 2 (P2F) and phase 3 (P3F). ‘ND’: Not detected indicated genera which were below the detection limit. a) Reactor Set B; received Cowansville Influent Solids b) Reactor Set C received Pincourt Influent Solids. μnet was modelled as described in section ‘Modelling net growth rate of bacteria’ and Table S3.

The remainder of the immigrants present at end of Phase 3 were classified as having a negative net growth rate (Figure 6). The detection of these taxa at the end of Phase 3 was unexpected, however it was noted that these genera typically occurred in higher abundance at the end of Phase 2. During Phase 3, the abundance typically reduced by at least 85 % suggesting that washout would likely occur over time. When excluding these taxa, an odds ratio test showed that taxa with a positive net growth rate were 14.0 to 15.2 times more likely to persist at the end of Phase 3 than those with a negative net growth rate. Other factors, such as the influent solids as an additional food source also likely influenced the persistence or loss of these taxa during Phase 3.

### Impact of Immigration on Core Resident Genera

The quantitative assessment of immigration reported thus far focused exclusively on immigration dependent communities, i.e., those which are introduced into the system and only present with continuous immigration. Core resident genera were present under all reactor conditions, both with and without immigration, and accounted for up to 76 % of sequencing reads (Table 1). It was also noted these genera were often detected within the influent wastewater, with differences in the ASVs observed in the influent solids and the reactors receiving only Syntho. It could be hypothesised that although the core resident population remains relatively stable over time in all reactors irrespective of feed received, immigration may contribute to diversity of these genera at ASV level.

Analysis of the ASV richness of core resident genera at the end of Phase 2 (Figure 7) showed a significant increase with immigration in Set A and Set B of the reactors (Figure 7a and 7b; unpaired t-test P = 0.008 and P= 0.02 respectively). A significant increase was not observed in Set C of the reactors (Figure 7c), where the impact of immigration on ASV diversity appeared to be more variable. In total between 44 and 73 unique ASVs belonging to genera classified as core residents were identified in the influent microbial community (Figure 7d), demonstrating that immigration had an impact on the core resident community at ASV level. However, between only 10 to 17 of these ASVs were detected in the reactors receiving influent solids, indicating selection to occur between these settings. Of those successfully immigrating were ASVs from the genera *Acinetobacter*, *Pseudomonas* and *Zoogloea*. The genus *Acinetobacter*, which was highly diverse without immigration (up to 15 ASVs) had 3 additional ASVs introduced. However, selection was also observed, and several *Acinetobacter* influent ASVs were not detected in the reactors. Multiple sequence alignment was used to study the sequence similarity between the ASVs (Figure S3a) and it was observed that ASVs which formed the core *Acinetobacter* population and successful immigrants typically clustered together.

**Figure 7.**
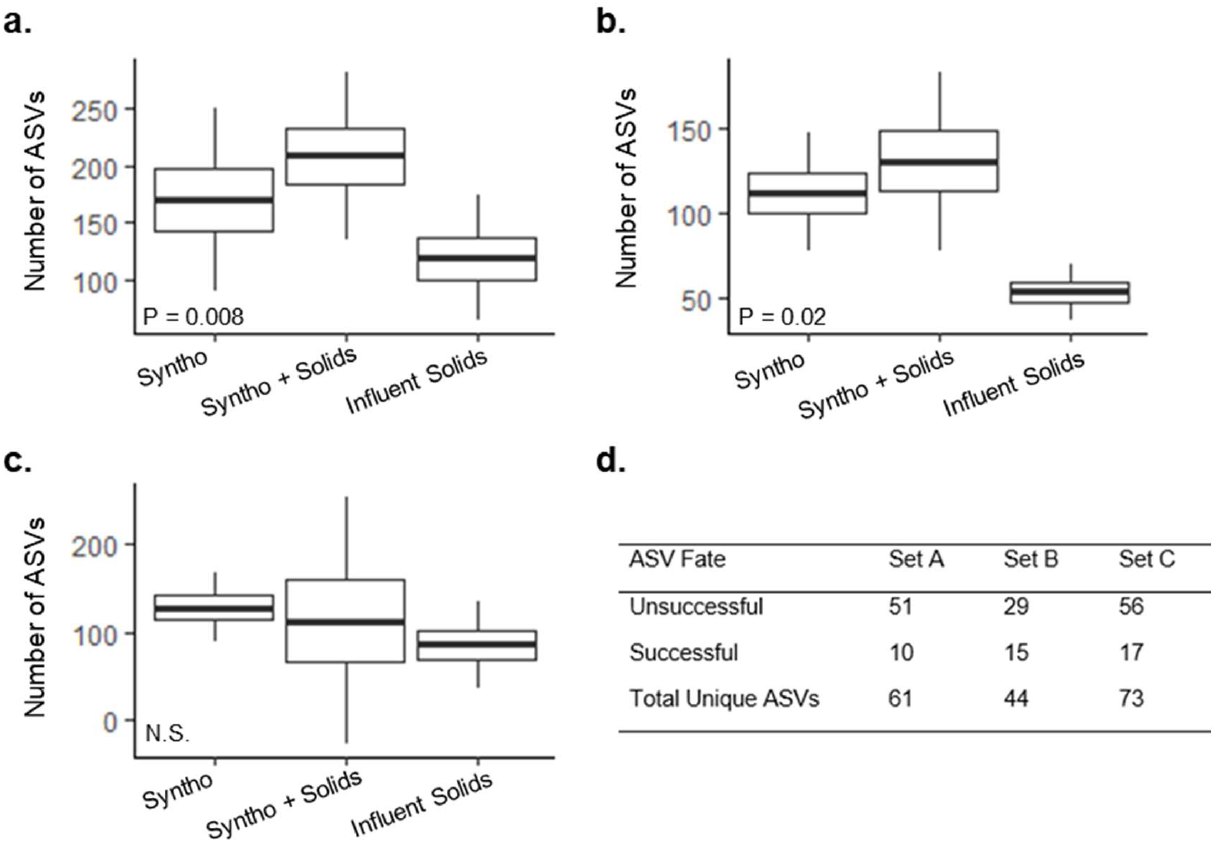
Impact of immigration on ASV richness of the core resident genera. a) Reactor Set A received La Prairie influent solids, b) Reactor Set B received Cowansville influent solids, c) Reactor Set C received Pincourt influent solids. d) The fate of unique influent ASVs in the reactors; ‘Unsuccessful’ were those detected in the influent, but not in the reactors, ‘Successful’ were those which were detected in the reactors with immigration.

Similarly, up to 4 additional *Pseudomonas* ASVs were introduced with immigration whilst other influent ASVs did not successfully immigrate. Multiple sequence alignment of the *Pseudomonas* ASVs did not show clear clustering based on the fate of a given ASV (Figure S3b). In the genus *Zoogloea* all influent ASVs were detected in the reactors with immigration. Multiple sequence analysis was performed to look at the sequence similarity between each ASV and to further investigate patterns of selection (Figure S3c). Among the *Zoogloea* ASVs, no clustering was observed based upon their occurrence in the core resident populations or influent solids.

## Discussion

The importance of influent immigration has long been debated among technical professionals, theorists, and modellers alike. One factor obscuring this debate in studies of full-scale wastewater treatment plants is the impossibility to differentiate the impact of influent substrate landscape from the immigration of microbial populations. To achieve this differentiation, the current study aimed to develop a reproducible and controlled system to investigate the impact of immigration and its relative contribution to the wastewater treatment plant community. To determine the fate of immigrating bacteria both substrate and continuity controls were included and to assess the impact of the AS-WWTP community itself, three different inoculums were used.

### Wastewater Substrate Composition and Operation Cause Communities to Become More Similar

The impact of wastewater substrate composition on the activated sludge community is well documented in the literature (34, 35). Thus, it was as expected that during Phase 1 the introduction of a common wastewater source and identical operational conditions caused the inoculum communities to become more similar (Figure 2). As discussed, comparing the Jaccard and Bray-Curtis dissimilarities of the communities at the end of Phase 1 revealed a higher difference measured by Jaccard dissimilarities (ANOSIM R = 0.88 + 0.05) than Bray-Curtis dissimilarity (ANOSIM R = 0.61 + 0.04; Figure 2). Taken together this suggested that the influent’s substrate composition selects for a core resident microbial community, whilst features of the inoculum were associated with lower abundance genera.

At the end of Phase 2, reactor Set A and B remained significantly clustered based upon the inoculum received (Figure S1 and Table S3; Jaccard distance, ANOSIM R= 0.77 + 0.13). However, in reactor Set C, clustering was not observed in the test reactors which received influent solids, suggesting that features of the starting community were impacted by immigration (Table S3).

On a larger scale, with an increase in the popularity of microbial transplantation research over recent years to improve feed efficiency in livestock (36), or to restore function of commensal gut microbiota and reduce the prevalence of multidrug resistant organisms (37), the plausibility of activated sludge bioaugmentation has been raised. This concept would involve taking a desired activated sludge microbial community from one location and transplanting it into a AS-WWTP requiring optimisation. The current study suggests activated sludge bioaugmentation may have variable results, as the activated sludge community is largely dependent on the wastewater substrate composition and not the inoculum alone.

### Immigration Impacted the Microbial Community of the Activated Sludge

Previous studies have inferred the impact of immigration based upon shared taxa between the influent and AS-WWTP, which provides limited information about the microbial activity of these organisms within the AS-WWTP. The inclusion of a sterile substrate control in this study allowed us to quantitively assess the impact of immigration, enabling the active and inactive portions of the immigrating community to be distinguished experimentally. Active immigration-dependent genera were defined as those only present in reactors receiving influent solids, and in a higher abundance than in the sterile control, indicating growth of the taxa. Actively growing immigration dependent taxa were found to account for between 4 to 14 % of the total reads in the mixed liquor, contributing a significant proportion of the community (Table 2). This result was consistent with previous studies that reported 10 % of OTUs to be present primarily due to immigration (9). Considering the significant proportion of activated sludge reads contributed through immigration, this source should not be neglected in process design and optimisation. Nonetheless, due to the low abundance of immigrant populations, analysis at ASV level was not possible. Future immigration studies should consider performing deeper sequencing strategies to enable analysis of ASV diversity in taxa occurring at low abundance.

Beyond immigration-dependent genera, immigration also impacted core resident genera, which were defined based upon their presence in at least 80 % of reactors under all conditions (with and without immigration) (Table 1). It was observed that genera classified as core residents were often present in the influent wastewater, and the ASV composition of these genera was different between the two environments. Analysis showed a significant increase in diversity of the core resident population with immigration in two of the three sets of reactors (reactor Set A and B; Figure 7a & 7b). Successful immigration was observed between the influent and mixed liquor with between 1 and 6 % of reads contributed by immigrating ASVs. As a result, the steady-state model used in this study may somewhat inaccurately estimate the growth rate of some core resident genera receiving immigrant ASVs.

Selection was an important factor filtering the immigrating populations (Figure 7d), as the majority of unique ASVs identified in the influent solids were not detected in the reactors with immigration. Overall, between only 16 and 34 % of ASVs unique to the influent wastewater successfully immigrated into the reactors. This is consistent with previous studies which show that heterotrophs are strongly selected between influent wastewater and the activated sludge (12). This could be due to niche differences between the environments, or competition for resources with the well-established, metabolically similar reactor core resident community. The influence of selection varied at genus level among the resident core genera (Figure S3). For example, ASV selection was observed in genus *Acinetobacter* but not in genus *Zoogloea*. This supports the proposal that heterotrophs should not be considered as a whole, and instead should be divided into relevant functional guilds for proper dynamic analyses.

### Growth Rate is a Key Determinant in the Fate of Immigrants

Analysis of the resident and immigrant populations (Figure 5) showed differences in the net growth rates. Resident genera typically exhibited positive net growth rates, explaining how they are present without constant seeding. Conversely, immigrant genera typically displayed low or negative net growth rates, particularly in reactor Set B and C. Based upon the terminology proposed by Frigon and Wells (2019) the majority of immigrants could be classified as mass-flow immigrants (i.e., continuous immigration at a high rate), as they displayed low or negative net growth rates in the mixed liquor (17). This could be attributed to differences in niche availability between the influent wastewater and the reactor communities. This result is consistent with that predicted by studies using a mass balance approach, which showed a large proportion of taxa shared between the influent and AS-WWTP to have a negative net growth rate (9). It was concluded that this fraction of the immigrant community was inactive and did not contribute to the metabolism of the activated sludge. However, despite these findings, we empirically demonstrated that although displaying a negative net growth rate, active immigration-dependent genera are in fact growing and consuming substrates within the activated sludge because their abundance was higher than in reactors receiving fresh solids than in reactors receiving autoclaved solids. This observation is in agreement with previous studies that demonstrated the rescue of complete nitrification in activated sludge bioreactors by nitrifiers in the influent stream (11). Consequently, observations of shared OTUs (ASVs or taxa) do not accurately predict activity, and more work should be done to quantify the metabolic contribution of immigrants with net negative growth rates.

During Phase 2, continuous influent immigration prevented the competitive exclusion of the immigrating bacteria, which were not well adapted to the reactor systems. This was confirmed during Phase 3 (reactor Set B and C only), when reactors received synthetic wastewater only and immigration was removed. During this phase, a large proportion of immigration-dependent genera were washed out within 3 SRTs (Table 3).

Of those remaining, between 75 and 77 % exhibited a positive net growth rate during Phase 2, which is consistent with reactor theory that these genera could be maintained within the reactors without the need for continuous immigration. However, other genera classified as actively growing immigration-dependent genera which displayed a positive net growth rate during Phase 2 (Figure 5) were not detected at the end of Phase 3. The discrepancy of these counter observations with reactor theory could be due to the imprecisions and inaccuracies in the quantification of growth rates. Nonetheless, they could also be explained by various abiotic and biotic phenomena. Abiotic phenomena include the possibility that influent solids provided an additional food sources or other beneficial niche markers for these genera which were removed during Phase 3. Biotic explanations may come from the co-selection of immigration-dependent populations as a single unit, but with some unit members exhibiting positive net-growth rates and others exhibiting negative net-growth rate (38). The reproducibility of the experimental system presented here would allow these different mechanisms to be explored in detail in future experiments.

The remainder of immigrants present at the end of Phase 3 displayed a negative overall net growth rate (Figure 6). Generally, these taxa had a higher relative abundance (0.1 % or above) at the end of Phase 2, whilst those with lower abundance were washed out by the end of Phase 3. A notable reduction (85 % and above) in the abundance of these genera was observed within 3 SRTs, suggesting that washout would likely occur with prolonged reactor operation and no immigration.

### Resident Core Genera

Detailed studies of the microbial community composition of AS-WWTP samples collected worldwide has enabled core taxa to be identified based upon their occurrence in different locations and their relative abundance (33). The core resident genera of the reactor communities overlapped with those previously reported in full-scale wastewater treatment plants. For example, some of the most abundant resident core genera such as *Zoogloea, Haliangium* and *Flavobacterium* were previously classified as strict core genera, which were observed in at least 80 % of the AS-WWTPs with a relative abundance >0.1 % (33). Whilst other abundant reactor core residents such as *Aeromonas* and *Thermomonas* were classified as ‘general core’ taxa, which occurred in 50 % of full-scale AS-WWTP at an abundance of >0.1 %. Taken together, these results demonstrate that the core community produced during this reactor study is representative of that in full-scale AS-WWTPs.

### Growing Immigrant Genera

The community of immigrating bacteria varied based upon the source of influent solids received. Among reactor Set A and B, the genera Spb280 and *Aquabacterium* respectively (both family *Comamonadaceae*) were dominant immigrants (Figure 4) and found exclusively within these reactor sets. Members of the *Comamonadaceae* family have previously been identified as habitat generalists capable of surviving in both influent wastewater and activated sludge (17). Thus, the presence of these immigrants is consistent with results from full-scale AS-WWTP immigration studies.

Among reactor Set C, which received Pincourt influent solids, the genera in the immigrant population were more variable (Figure 4). Whilst some abundant genera such as *Dechloromonas* appeared in all reactors, others such as *Elstera* and *Candidatus* Accumulimonas appeared more sporadically. This could be due to differences in niche availability in reactors or co-selection. Limitations associated with small-scale reactor experiments should also be considered. Given the influent solid particle size and the small reactor volume, there may have been a slight variability in the microbial community of the influent solids received by each reactor that exacerbated the observed community drifts.

Among the reactors, few examples of inoculum effect were observed. This validates this reactor protocol to be suitable for future immigration (or perpetual coalescence) studies, regardless of the mixed liquor inoculum source. In reactor Set B, one example where inoculum appeared to have an impact, was in the genus SipK9 (family *Xanthomonadaceae)*, which was dominant only among reactors receiving inoculum A (La Prairie) and accounted for between 2.5-11.7 % of overall reads. SipK9 is an uncultured genus which has been previously reported in Antarctic mineral soils (39) and activated sludge AS-WWTPs as documented in the MiDAS ecosystem specific reference database (33). In the reactors inoculated with Pincourt and Cowansville ML, the maximum abundance of SipK9 was 0.008 % indicating that there were differences in niche availability or co-selection between the reactors based on the inoculum received. The function of this population is currently unknown. However, its high abundance warrants further investigation.

Control over specific populations may be key to both process optimisation and mitigation of operational problems in activated sludge. In the immigrating community specialised bacteria such as the phosphorus accumulating organisms *Dechloromonas*, *Candidatus* Accumulimonas, *Candidatus* Accumulibacter and *Tetrasphera* were identified (Figure 4). Given the importance of these genera in enhanced biological phosphorous removal, immigration may be key in process optimisation. Unfavourable taxa such as *Gordonia*, a filamentous bacteria, and *Thauera*, which cause bulking and dewatering problems in activated sludge treatment plants, were also identified (40). By accounting for the degree of immigration in the design of solutions, operational problems in AS-WWTPs could be managed more efficiently.

## Acknowledgments

We would like to thank Julia Qi, Carlos Vasquez Ochoa, Nouha Klai and Zeinab Bakhshijooybari for their assistance with reactor operation. We are also indebted to the operators and staff at Cowansville, La Prairie and Pincourt wastewater treatment plants for access to the facilities and help with sampling.

This work was funded by NSERC through a Discovery grant (NSERC RGPIN-2016-06498) and a Strategic Program Grant (NSERC STPGP 521349-18). NSERC had no role in the design of this study.

Claire Gibson, Shameem Jauffur, and Bing Guo were partly funded by the McGill University Engineering Doctoral Award.

