## Supplementary Information for "Activated Sludge Microbial Community Assembly: The Role of Influent Microbial Community Immigration"

* Present address: Environmental Department, Amazon Web Services, Canada

| Table S1- Phases of reactor operation | | | | |
| --- | --- | --- | --- | --- |
| Phase | Reactor Feed |  | Duration (SRTs) ^1^ | |
|  |  |  | 2016 | 2017 |
| 1 | All receive Syntho only |  | 12 | 10 |
| 2 | Reactors receive three different feeds^2^ |  | 13 | 22 |
| 3 | All receive Syntho Only |  | NA^3^ | 3 |
| ^1^ One SRT represents 5 days of reactor operation  ^2^ Three feeds; Syntho and influent solids, Syntho and sterile solids and Syntho Only  ^3^ NA: not available; reactors in 2016 were not operated with a third phase | | | | |

| Table S2: Overview of the Distribution of Reactors in the Experimental Design (**Total Reactors:** 72) | | | | | | | | | | | | |
| --- | --- | --- | --- | --- | --- | --- | --- | --- | --- | --- | --- | --- |
|  |  | **Syntho + Influent Solids** | | |  | **Syntho + Sterile Solids** | | |  | **Syntho Only** | | |
|  |  | Block^1^ | | |  | Block | | |  | Block | | |
| Set^2^ |  | **a** | **b** | **c** |  | **a** | **b** | **c** |  | **a** | **b** | **c** |
| **A** |  | 3 | 3 | 3 |  | 3 | 3 | 3 |  | 3 | 3 | 3 |
| **B** |  | 3 | 3 | 3 |  | 3 | 3 | 3 |  | 3 | 3 | 3 |
| **C** |  | 3 | 3 | 3 |  | 3 | 3 | 3 |  | Same as Set B^3^ | | |
| ^1^ Where Block a, b and c represent the different sources of the inoculum  ^2^ Where Set A, B and C represent the different sources of influent solids  ^3^ Syntho Only Control was shared for set B and C as the experiments were conducted at the same time | | | | | | | | | | | | |

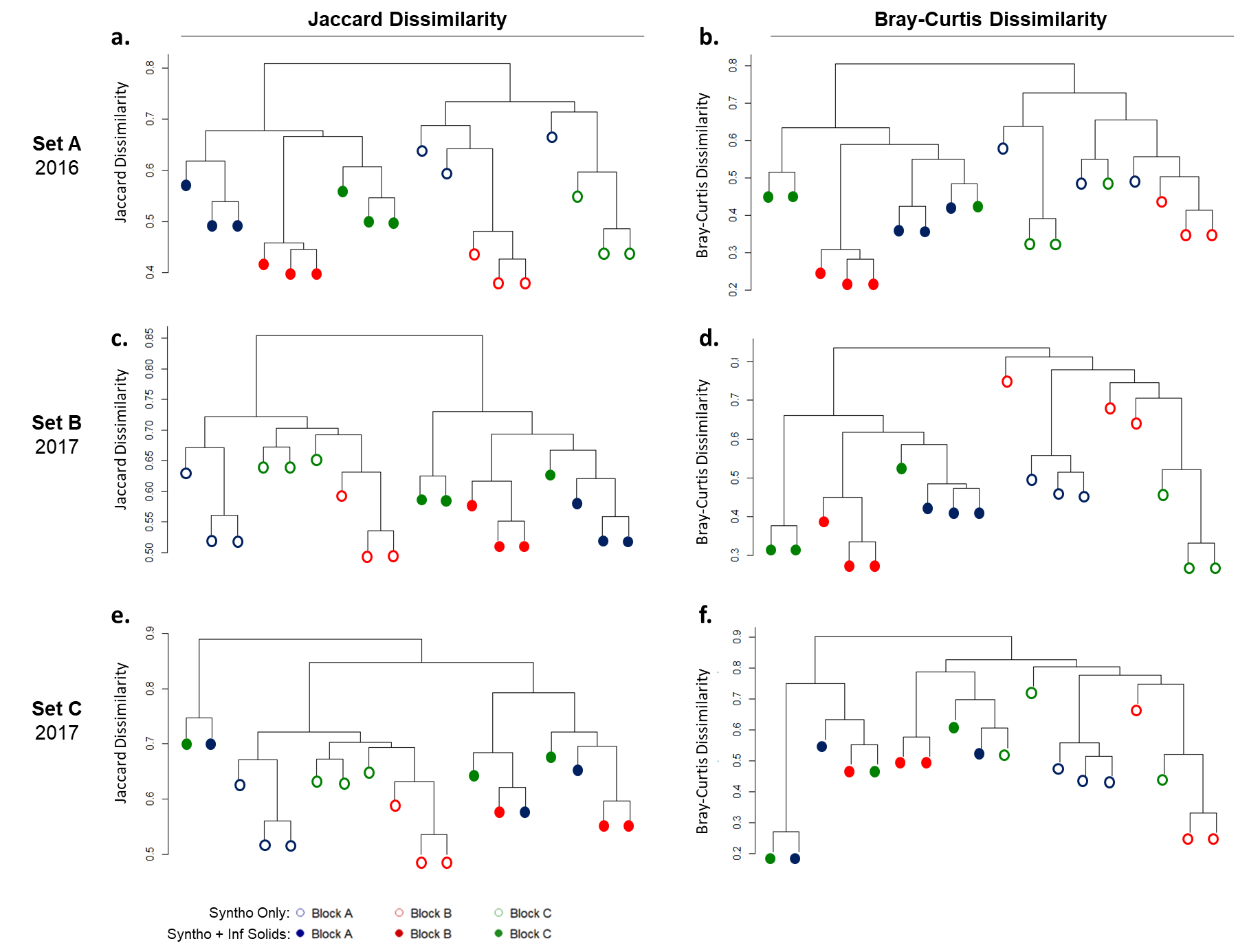

Figure S1- Tree dendrogram of microbial community at the end of Phase 2 using UPGMA clustering method and the Jaccard dissimilarity (a, c, e) or the Bray-Curtis dissimilarity (b, d, f). Filled symbols indicate test reactors which received influent solids during Phase 2. Hollow symbols indicate samples which received Syntho only throughout. Set A (a, b) were operated in 2016, whilst Sets B (c, d) and Set C (e, f) were operated in 2017. The Block indicates the source of inoculum: Block a-La Prairie mixed liquor, Block b-Cowansville mixed liquor, Block c-Pincourt mixed liquor. Inoculum communities were the same for Set B and C as they were operated in tandem.

| Table S3- ANOSIM Analysis of Microbial Community at the End of Phase 2 (Figure S1) | | | | | | |
| --- | --- | --- | --- | --- | --- | --- |
|  |  | Jaccard Dissimilarity | |  | Bray-Curtis Dissimilarity | |
|  |  | ANOSIM R | α^1^ |  | ANOSIM R | α |
| **Set A** | Syntho + Influent Solids | 0.96 | 0.005 |  | 0.84 | 0.008 |
|  | Syntho Only | 0.61 | 0.013 |  | 0.53 | 0.017 |
| **Set B** | Syntho + Influent Solids | 0.75 | 0.003 |  | 0.75 | 0.007 |
|  | Syntho Only | 0.77 | 0.001 |  | 0.46 | 0.02 |
| **Set C** | Syntho + Influent Solids | NS^2^ | NS |  | NS | NS |
|  | Syntho Only | 0.77 | 0.003 |  | 0.46 | 0.017 |
| ^1^ α – Significance level  ^2^ NS- Not significant | | | | | | |

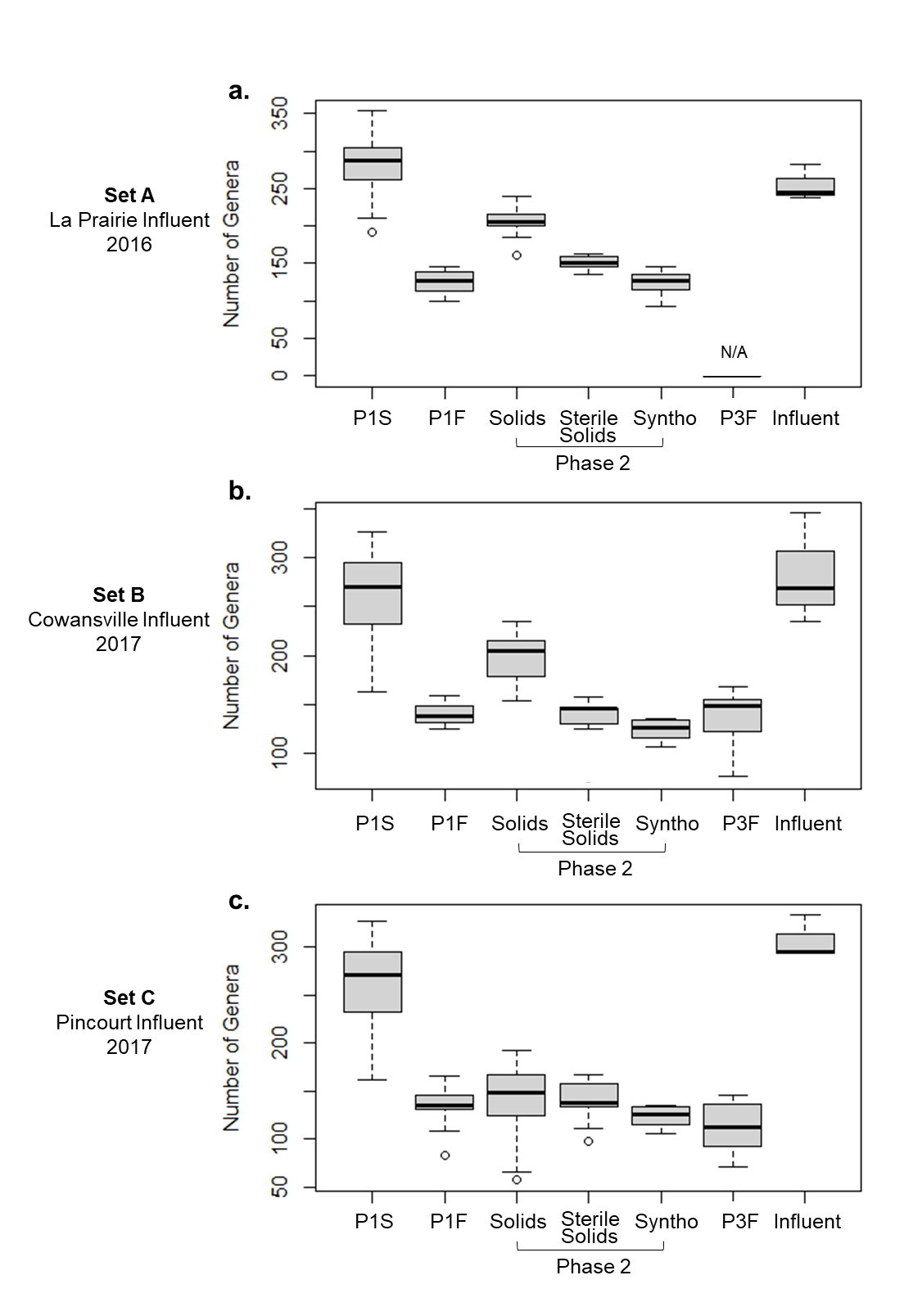

Figure S2- Genus richness boxplot of reactor communities in the inoculum at the beginning of the experiment (P1S; n = 9), at the end of Phase 1 (P1F; n = 27), end of Phase 2 (Solids: fed Syntho + influent solids, Sterile Solids: Syntho + autoclaved influent solids, and Syntho only; n = 9 each), end of Phase 3 for reactors receiving Syntho + influent solids (P3F; n = 9), and Influent Solids (n = 3). Phase 3 was completed for only reactor Sets A and B. a) Set A received La Prairie influent solids during Phase 2 b) Set B- Cowansville influent solids received c) Set C- Pincourt influent solids received. In the boxplot the middle bar represents the median number of genera within sample group. Box edges represent the first and third quartile. Whiskers represent the minimum and maximum number of genera.

| Table S4: Values used in the calculation of the relationship between $f_{16S,i,ML}, f_{16S,i,inf}$, and $\mu_{net,i}$ | | | | |
| --- | --- | --- | --- | --- |
| **Parameter** | **Symbol** | **Reactor Set** | | |
|  |  | A- La Prairie | B- Cowansville | C- Pincourt |
| Solid Retention Time (days) | $\theta_{x}$ | 5.0 | 5.0 | 5.0 |
| Hydraulic Retention Time (days) | $\theta$ | 1.8 ± 0.28 | 1.8 ± 0.28 | 1.8 ± 0.28 |
| Fraction of influent captured by ML solids | $f_{OHO,Capt}$ | 1 | 1 | 1 |
| Total VSS of the Influent (mg-VSS/L) | $X_{Tot,Inf}$ | 120 | 120 | 120 |
| Total VSS of the ML (mg-VSS/L)^1^ | $X_{Tot,ML}$ | 470.9^2^ | 685.3 ± 94.22 | 644.4 ± 73.54 |
| DNA extraction yield of Influent (µg-DNA/mg-VSS)^3^ | $\gamma_{DNA,Inf}$ | 1.73 ± 0.15 | 0.72 ± 0.07 | 0.94 ± 0.13 |
| DNA extraction yield of ML (µg-DNA/mg-VSS)^4^ | $\gamma_{DNA,ML}$ | 1.51 ± 0.09 | 1.75 ± 0.004 | 1.06 ± 0.09 |
| ^1^Where n=18 representing the average VSS of the nine reactors receiving influent solids (with immigration) over a two week period | | | | |
| ^2^Calculated based upon COD/VSS ratio and validated with experimental data obtained during Phase 1 as no experimental measurements | | | | |
| were obtained for this reactor set during Phase 2 | | | | |
| ^3^Influent DNA yield where n=3 and samples taken from three different time points | | | | |
| ^4^Mixed liquor DNA yield where n=3 for each reactor set (1 reactor per block within each set). Samples obtained from the final day of operation | | | | |
| ± indicates the standard deviation of reported measurements | | | | |

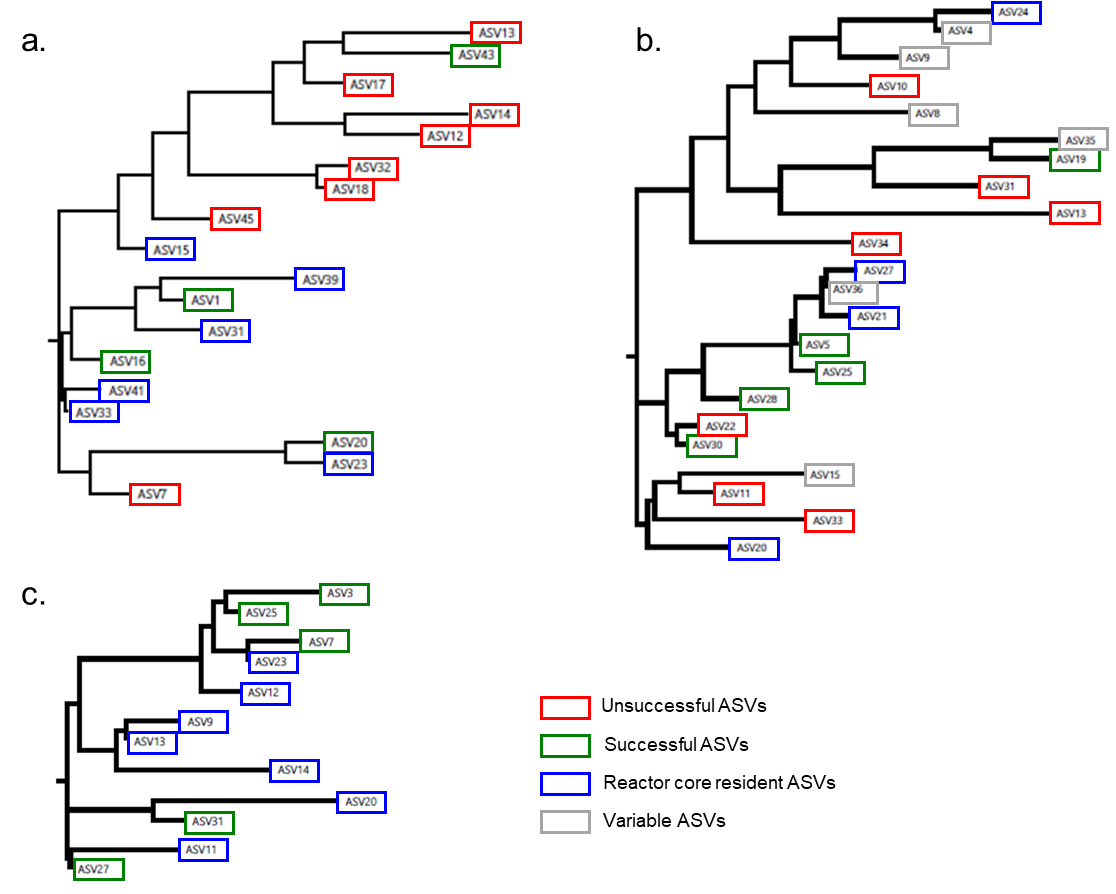

Figure S3- Multiple sequence alignment tree generated using BIONJ neighbour joining algorithm. a) Acinetobacter ASVs b) Pseudomonas ASVs c) Zoogloea ASVs. Unsuccessful ASVs are those that were detected in the influent but not in reactors with immigration. Successful ASVs are those detected in the influent and in the reactors with immigration only. Reactor core resident ASVs were those present in all reactors regardless of feed. Variable ASVs are those which had different fates in each reactor set (e.g. successful in one reactor set and unsuccessful in another).
